# Sashimi.py: a flexible toolkit for combinatorial analysis of genomic data

**DOI:** 10.1101/2022.11.02.514803

**Authors:** Yiming Zhang, Ran Zhou, Yuan Wang

**Author notes:** Correspondence (R.Z.), (Y.W.). These authors contributed equally to this work (Yiming Zhang, Ran Zhou).

## Abstract

Simultaneously visualizing how isoform expression, protein–DNA/RNA interactions, accessibility, and architecture of chromatin differs across condition and cell types could inform our understanding on regulatory mechanisms and functional consequences of alternative splicing. However, the existing versions of tools generating sashimi plots remain inflexible, complicated, and user-unfriendly for integrating data sources from multiple bioinformatic formats or various genomics assays. Thus, a more scalable visualization tool is necessary to broaden the scope of sashimi plots. Here, we introduce sashimi.py, a Python package for generating publication-quality visualization by a programmable and interactive web-based approach. Sashimi.py is a platform to visually interpret genomic data from a large variety of data sources including single-cell RNA-seq, protein–DNA/RNA interactions, long-reads sequencing data, and Hi-C data without any preprocessing, and also offers a broad degree of flexibility for formats of output files that satisfy the requirements of major journals. The Sashimi.py package is an open-source software which is freely available on Bioconda (https://anaconda.org/bioconda/sashimi-py), Docker, PyPI (https://pypi.org/project/sashimi.py/) and GitHub (https://github.com/ygidtu/sashimi.py), and a built-in web server for local deployment is also provided.

## 1 Introduction

Differential isoform expression plays a critical role in the diversification of the proteome and functionality of transcripts [1]. A variety of library protocols or sequencing methods has been established and become widely used to address this problem such as single-cell RNA-seq (scRNA-seq) [2] or long-read sequencing [3]. Though multiple advanced tools have been developed, the analysis and visualization of these informative genomics data still face several challenges. On the one hand, popular tools including sashimi [4], ggsashimi [5], and SplicePlot [6] lack support for a wide range of bioinformatic format and extra preprocessing is necessary for complex data such as demultiplexing scRNA-seq or scATAC-seq by cell barcodes. Moreover, interactive genome browsers like Integrative Genomics Viewer (IGV) [7] requires time-consuming loading of voluminous alignment files. On the other hand, most of the existing tools only offer a command-line interface to use which is unfriendly to inexperienced programmers. To address these limitations, we present Sashimi.py, a new and comprehensive tool to generate publication-quality plots in a programmable and interactive web-based approach. Sashimi.py supports the integrated visualization for a large variety of data sources, such as gene annotation with functional domain mapping, isoform expression, isoform structures revealed by scRNA-seq and long-read sequencing, as well as accessibility, and architecture of chromatin.

## 2 Description

Sashimi.py is built by python and JavaScript which greatly facilitates future maintenance. It’s a platform to visualize genomic data from a variety of sources on the given genomic coordinate and generate an image including all of the input datasets. It supports most various standard data formats in bioinformatics, including BAM, BED, bigWig, bigBed, GTF, BedGraph, HiCExplorer’s native h5 format, and the depth file which is generated by samtools [8] (Figure 1). Sashimi.py is also an easy-access package with highly reproducible, and it could be freely downloaded from GitHub, and installed from source code, PiPY, Pipenv, and Bioconda as well as a Docker image (Figure 1).

**Figure 1.**
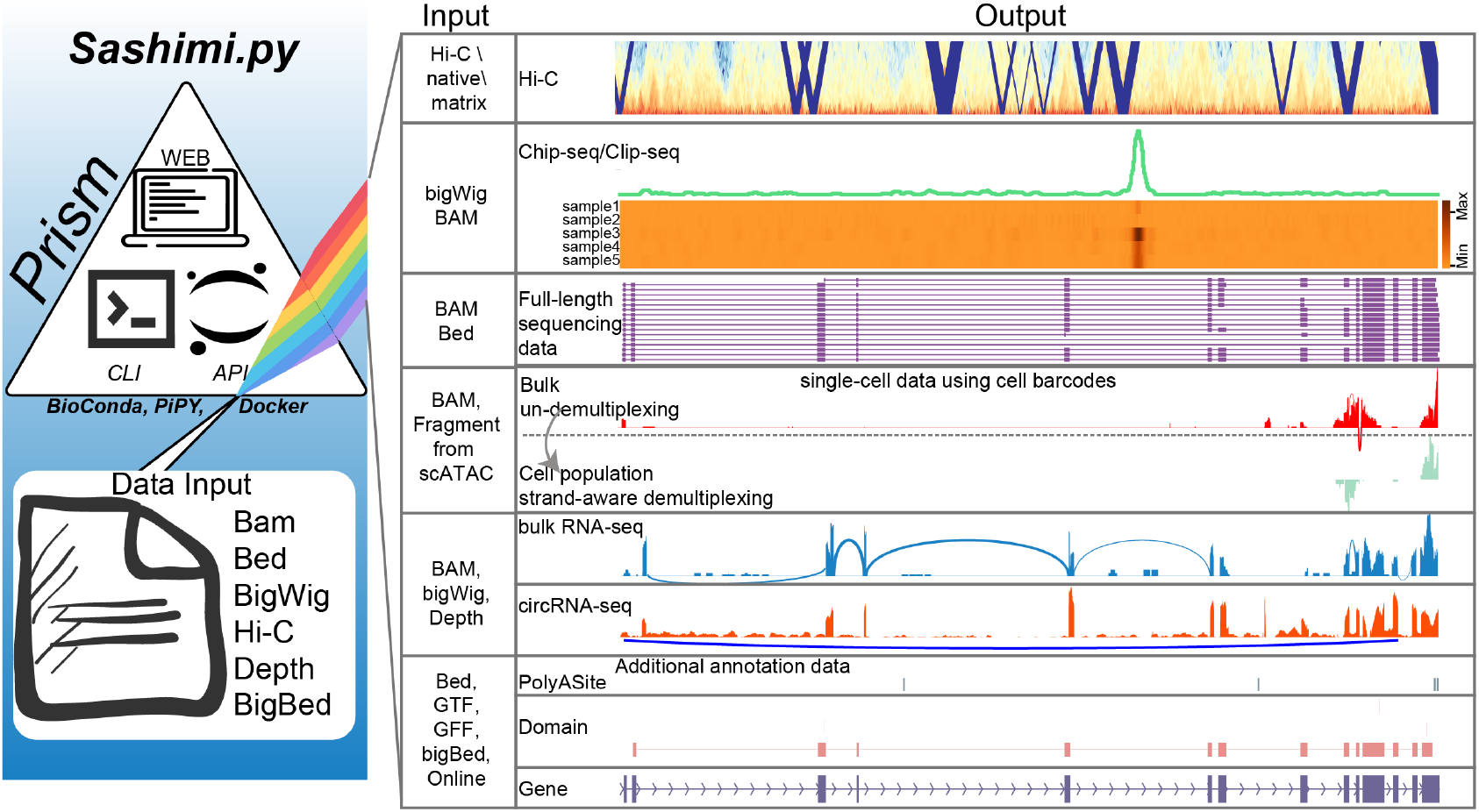
The schematic diagram of the Sashimi.py tool. Sashimi.py, which is built by python and JavaScript that greatly facilitates future maintenance, could visualize genomic data from a variety of sources and bioinformatic formats on the given genomic coordinates.

Sashimi.py offers a variety of approaches to use, and users could generate the desired plots by an application programming interface (API) from a script or Jupyter Notebook as well as a command-line interface (CLI). In API mode, users could configure and generate plots from multiple genomic data without any preprocessing in a block building manner. This greatly facilitates users for integrating Sashimi.py into scalable pipelines or analysis frameworks. Moreover, Sashimi.py also offers a local web application implemented in JavaScript, and users could rapidly deploy the local webserver to render the plot of the desired tracks in a mouse-click. Besides, The CLI with the detailed introduction is also provided by Sashimi.py, and the only preprocessing step is to prepare the configuration file which contains all necessary parameters for each track. Furthermore, a detail-oriented online document (https://sashimi.readthedocs.io/en/latest/) also is provided by Sashimi.py which includes all examples of the different approaches.

For plot generation, the precise genomic coordinate of interest is required by Sashimi.py. It also offers a broad degree of flexibility for the size and resolution (dots per inch, DPI) of the figure as well as output formats such as *png, pdf* and *tiff* which satisfy the requirement of the major scientific journals. Therefore, we present a highly accessible, reproducible and flexible tools for generating genomics data plots.

## 3 Use Cases

### Case1: Integration of multiple NGS data sources for revealing splicing regulation and outcomes

Same as previous sashimi plot packages, Sashimi.py takes splicing reads as input from BAM file and gene model annotation in GTF or BED file to visualize differential usage of exons or transcripts. An example picture for eight bulk RNA-seq samples from TNP GBM model [9] is shown in Figure 2A. The samples are colored by the stage of each sample and coverage densities across exons are calculated by aggregating reads. The plot highlights the differential splicing of the middle exon, which appears to be predominantly included at the T1 stage but mostly excluded at the end stage during the tumorigenesis. Alternative splice (AS) of mRNA generates more protein isoforms from a single gene, thereby contributing to protein diversity. But previous sashimi plot tools couldn’t display straightly the alteration of protein domains due to AS regulation. Therefore, in addition to present distinct usage of the exon across conditions, Sashimi.py also maps the protein coordinate into genomic coordinate after automatically crawling the domain information from UniProt [10] and ENSEMBL [11]. It shows that the long isoform which encodes a protein with key functional domains is gradually spliced out, and the short isoform without the key domains becomes a major isoform during the tumorigenesis. For facilitating users with a poor network, Sashimi.py supports local domain files as input which are downloaded from UCSC in advance.

**Figure 2.**
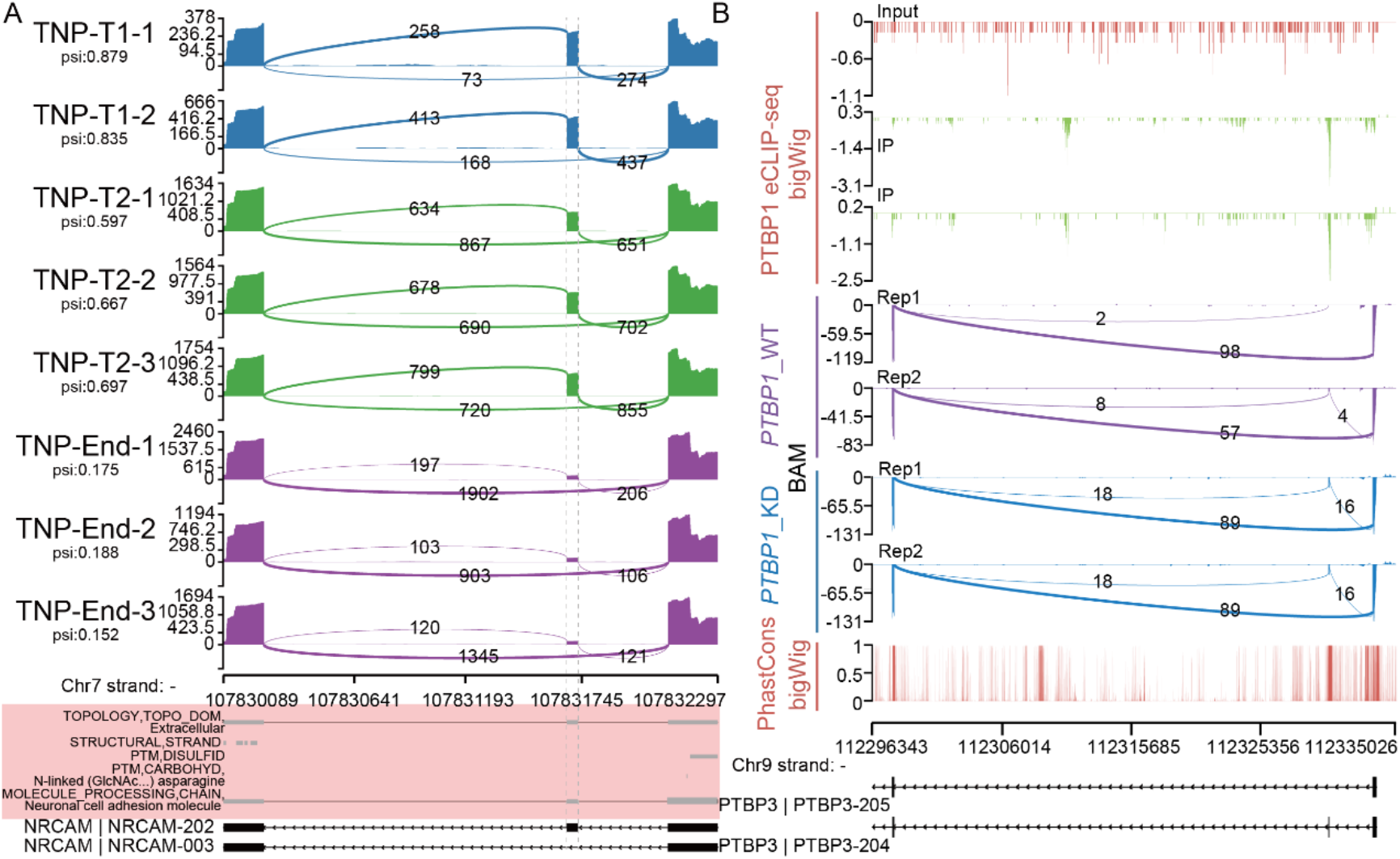
Integration of multiple NGS data sources for revealing splicing regulation and outcomes. (A) Sashimi plot with protein functional description for the time-course RNA-seq data with biological replicates from TNP model during tumorigenesis. Tracks indicated RNA-seq density. The span junction line indicated the middle exon (highlighted with a dash line) was decreasing during tumorigenesis, and its splice percent in (*?*) was highlighted below the label of each track. Bottom, the ENSEMBL gene annotation (black) and its protein domain information which was highlighted with pink background indicated different usage of isoforms with the distinct functional domains during tumorigenesis. (B) Sashimi plot for combinatorial isoform expression, RBP-RNA interactions and an evolutionarily conserved score of *PTBP3*. Tracks indicated the signal from different format files as input, including PTBP1 eCLIP-seq (bigWig), KD-RNA-seq (BAM) and PhastCons score (bigWig). Tracks for eCLIP-seq indicated that *PTBP1* directly binded the PTBP3 exon 2 with a highly conserved score (PhastCons track), and the exon 2 was significantly increased under perturbation of *PTBP1* (*PTBP1*_KD and *PTBP1*_WT tracks). Sashimi.py provides direct evidence that the alternative splice of PTBP3 exon 2 is likely regulated by PTBP1.

RNA binding proteins (RBPs) interact with RNA which governs the maturation and fate of the target RNA, and crosslinking and immunoprecipitation (IP) followed by sequencing (CLIP-seq) directly provides the set of candidate functional RNA element of the RBPs [12]. However, RBP binding signal is usually stored in bigWig format after normalizing and the existing sashimi tools couldn’t integrate it with splicing reads due to incompatible formats. With the ability to take multiple bioinformatic formats as input, Sashimi.py provides a flexible way to integrate data from multiple sources. As shown in Figure 2B, exon 2 of *PTBP3* was absent in the control cell but increased usage upon knockdown of *PTBP1*, consistent with previous findings [13]. And we observed strong biding events at splicing site of *PTBP3* exon 2 (Figure 2B), suggesting that this alternative event is likely to be directly regulated by PTBP3. Thus, Sashimi.py will facilitate users to integrate data from multiple sources and gain novel insight into the mechanism of RNA splicing across multiple conditions. In summary, Sashimi.py provides a highly flexible and compatible approach to reveal the potential function and regulation mechanism of AS events.

### Case2: Visualization of full-length sequencing data

The long reads sequencing (LRS) platform, such as Pacific Biosciences and Oxford Nanopore Technologies, provides the full structure of individual transcripts which do not require assembly. However, Current sashimi plot tools, which are designed for short-read sequencing data, visualize sequencing reads by aggregating depth of each coordinate and lost the connection of exons from the individual read. As shown in Figure 3A (top track), we observed a differential usage between exon 6 and exon 7 of *U2AF1* from the conventional sashimi plot on cerebral organoid data [14], but the advantage of LRS data which directly provides the full structure of transcript is not properly utilized. To maximize the advantage of LRS, Sashimi.py offers a read-by-read style with exon sort options that could distinctly present the exon-intron structures of each isoform. The exon 6 and exon 7 of U2AF1 represent a sub-category AS event–mutually exclusive exons (MXE), as we observed most of transcripts with exon 6 or exon 7 are likely original from *U2AF-201* or *U2AF-202* which encode a different protein (Figure 3A, bottom track), respectively. Moreover, data generated by Nanopore with a specific protocol also provides RNA modifications at single-nucleotide resolution such as N6-methyladenosine (m^6^A) [15]. In addition to present isoform structure, Sashimi.py could extract and visualize the extra information such as the length of poly(A) or modification status of each nucleobase from the tag of BAM file. As we observe most of the m6A sites are located in the last exon of *ACTB*, consistent with previous research [16] that the m6A sites are enriched in 3’ UTR. In summary, Sashimi.py provides a comprehensive approach to explore and visualize the whole structure and epigenetic modification of transcripts from LRS data.

**Figure 3.**
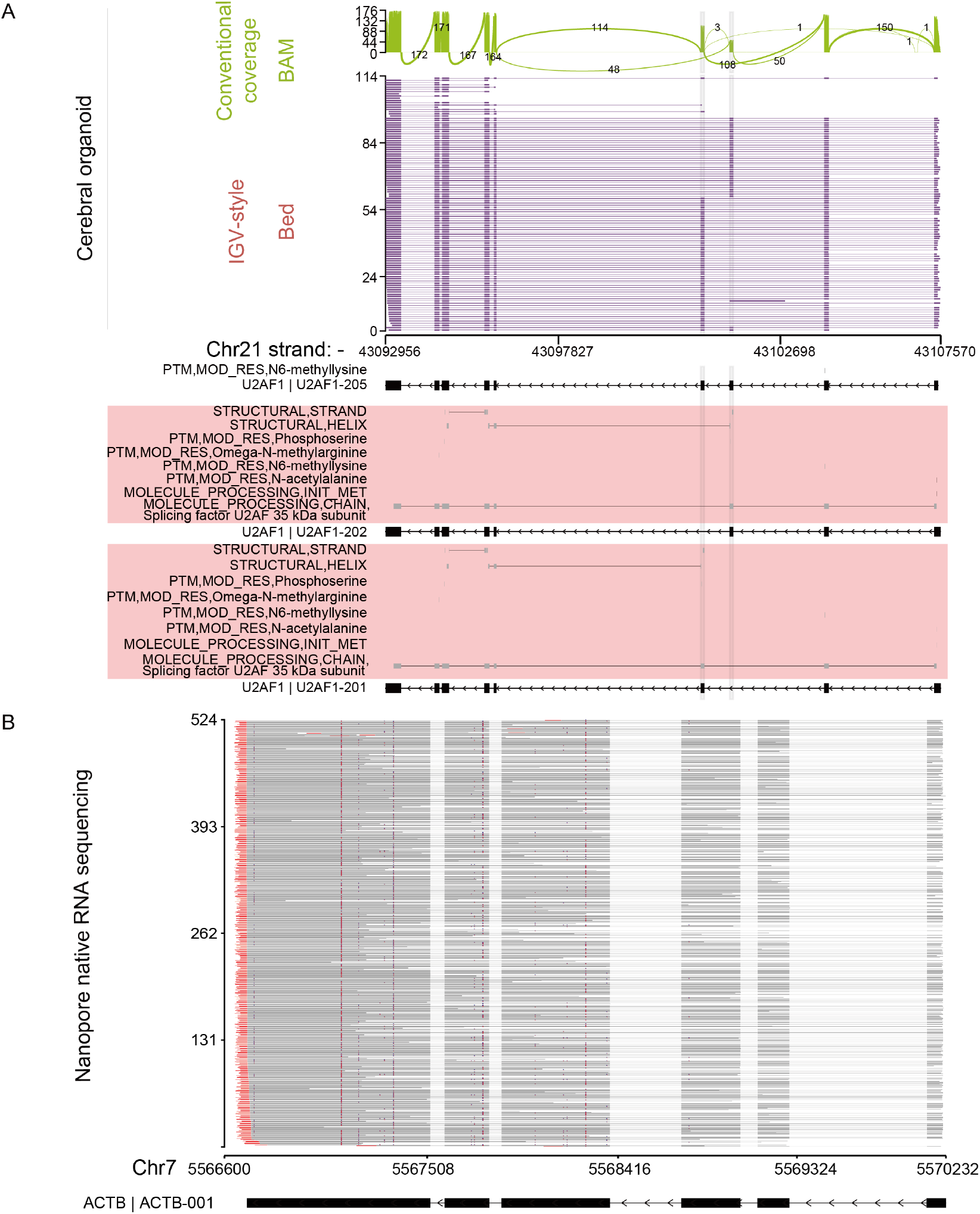
Visualization of full-length sequencing data. (A) Sashimi plot of cerebral organoid full-length sequencing data. The same data with different formats was presented by the conventional coverage plot and read-by-read plot that each line represented an individual read, respectively. The box highlighted a mutually exclusive exons (MXE) event. Bottom, the ENSEMBL gene annotation (black) and its protein domain information which was highlighted with pink background indicated the different functional protein outcomes caused by MXE. (B) Read-by-read track plot of nanopore native RNA sequencing data. The read part at end of each read and the blue dot on each read represented the length of poly(A) and the status of m6A modification.

### Case3: Decoding single-cell multiomics data

Recently, several methods such as SCAPE [17], which utilized the technologies inherently of 3’ based scRNA-seq, have been proposed to identify and estimate alternative polyadenylation events at single-cell level. However, due to the lack of the ability to demultiplex gene expression into cell population for the existing tools, users must manually split and deduplicate the bam file into individual files before subsequent processing. Fortunately, Sashimi.py could automatically demultiplex and deduplicate read depth based on the user-supplied meta file, which contains the cell barcode and its corresponding cell type. As shown in Figure 4A, three peaks which correspond to three distinct isoforms were identified in 3’UTR of *Tubb5-201* before demultiplexing (top panel). After strand-aware demultiplexing by Sashimi.py, it suggests that only pA1 and pA3 were generated by alteration of 3’ processing of the gene *Tubb5-201*, as we observed two sense peaks on 3’UTR of *Tubb5-201*. Surprisingly, the middle peak showed an opposite stand suggesting that pA2 is original from an anti-sense isoform of the *Tubb5-201*. Therefore, Sashimi.py provides a clearer picture of differential expression APA events among 3’ enriched scRNA-seq data, which facilitates researchers to turn genome-wide insights into validation experiments.

**Figure 4.**
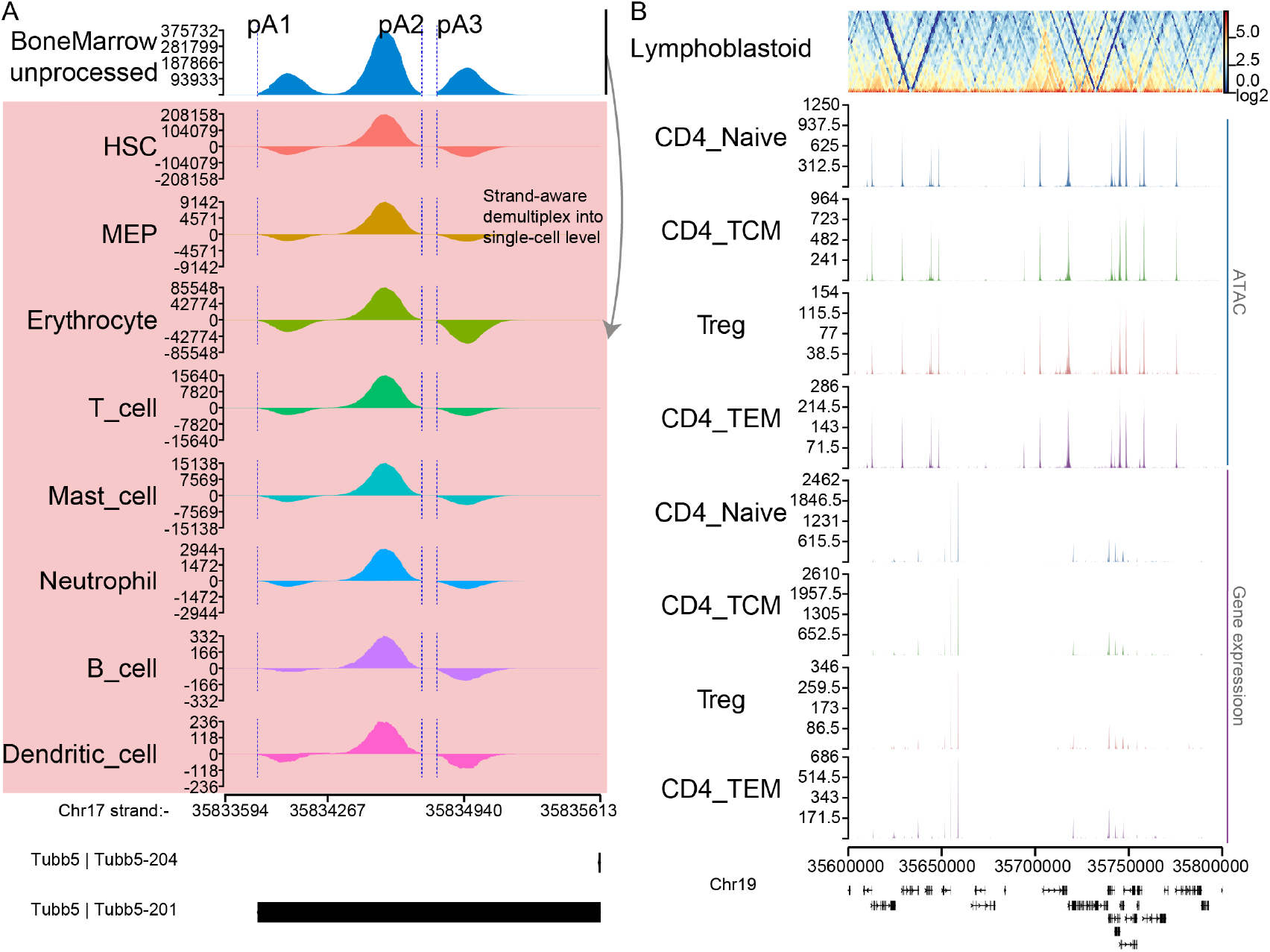
Decoding single-cell multiomics data. (A) Sashimi plot of 10x bone marrow datasets. Top tacks showed a density plot before demultiplexing, and tracks highlighted with pink background indicated strand-aware demultiplexing after Sashimi.py. The three peaks at each track represented three distinct isoforms and the polyadenylation site was indicated by a blue dash line which was identified by SCAPE. Bottom, the ENSEMBL gene annotation of *Tubb5*. (B) integration of single-cell multiomics and Hi-C data. The top track showed chromatin interactions from lymphoblastoid Hi-C data, and the color indicated the intensity of the interaction. The tracks labeled with a blue line and purple line represented the signal from simultaneous profiling of ATAC and gene expression for a cell, respectively.

Besides, the integration of transcriptome, chromatin accessibility, and chromatin architecture data provide a comprehensive understanding of functional enhancers regulating gene expression under different cellular or disease processes [18]. However, the existing tools couldn’t decode these multiomics data for visualization, especially single-cell transcriptome and chromatin accessibility data with cell barcodes. To end this, Sashimi.py also supports decoding single-cell data that simultaneously profile transcriptome and chromatin accessibility as well as chromatin interactions (Hi-C matrix) without any preprocessing. As shown in Figure 4B, it’s clear to observe the architecture and accessibility of chromatin as well as the gene expression pattern of lymphocyte cells on the interest genomic coordinate. In general, Sashimi.py provide a more comprehensive perspective to explore isoform diversity in cell population and potential enhancers for regulating gene and isoform expression.

## 3 Conclusion

With Sashimi.py, it is possible to integrate multiple data sources from a wide variety of genomic assays and generate publication-ready plots. It allows users to visualize NGS data with flexible formats of input files. To ensure maximal reproducibility, Sashimi.py also is distributed by PyPI, Bioconda and Docker. Sashimi.py also offers a flexible output format such as *png, pdf* and *tiff* which meets the requirement of major scientific journals. Sashimi.py can be used via a command line or an API for an interactive environment such as Jupyter Notebook. Moreover, it also provides a web application and users could deploy it on a local machine which provides an easy way to complete their analysis in a mouse-click. In summary, Sashimi.py offers an easy, fast, and reliable method for visualizing genomic data. Sashimi.py was written in Python and JavaScript which greatly facilitates future maintenance, and we will continue to maintain the updates and upgrades of the package based on suggestions and comments from the community.

## Data availability

The python package Sashimi.py is open-source and freely available on Docker, GitHub (https://github.com/ygidtu/sashimi.py), PyPI (https://pypi.org/project/sashimi.py/) and Bioconda (https://anaconda.org/bioconda/sashimi-py). The script to generate Figures 2-4 and several reproducible examples are available on GitHub (https://github.com/ygidtu/sashimi.py/example/Article_figures).

## Reference

1. Wright CJ, Smith CWJ, Jiggins CD. Alternative splicing as a source of phenotypic diversity, Nat Rev Genet 2022.

2. Macosko EZ, Basu A, Satija R et al. Highly Parallel Genome-wide Expression Profiling of Individual Cells Using Nanoliter Droplets, Cell 2015;161:1202–1214.

3. Jain M, Olsen HE, Paten B et al. The Oxford Nanopore MinION: delivery of nanopore sequencing to the genomics community, Genome Biol 2016;17:239.

4. Katz Y, Wang ET, Silterra J et al. Quantitative visualization of alternative exon expression from RNA-seq data, Bioinformatics 2015;31:2400–2402.

5. Garrido-Martin D, Palumbo E, Guigo R et al. ggsashimi: Sashimi plot revised for browser- and annotation-independent splicing visualization, PLoS Comput Biol 2018;14:e1006360.

6. Wu E, Nance T, Montgomery SB. SplicePlot: a utility for visualizing splicing quantitative trait loci, Bioinformatics 2014;30:1025–1026.

7. Thorvaldsdottir H, Robinson JT, Mesirov JP. Integrative Genomics Viewer (IGV): high-performance genomics data visualization and exploration, Brief Bioinform 2013;14:178–192.

8. Li H, Handsaker B, Wysoker A et al. The Sequence Alignment/Map format and SAMtools, Bioinformatics 2009;25:2078–2079.

9. Wang X, Zhou R, Xiong Y et al. Sequential fate-switches in stem-like cells drive the tumorigenic trajectory from human neural stem cells to malignant glioma, Cell Res 2021;31:684–702.

10. UniProt Consortium T. UniProt: the universal protein knowledgebase, Nucleic Acids Res 2018;46:2699.

11. Cunningham F, Allen JE, Allen J et al. Ensembl 2022, Nucleic Acids Res 2022;50:D988–D995.

12. Van Nostrand EL, Freese P, Pratt GA et al. A large-scale binding and functional map of human RNA-binding proteins, Nature 2020;583:711–719.

13. Tan LY, Whitfield P, Llorian M et al. Generation of functionally distinct isoforms of PTBP3 by alternative splicing and translation initiation, Nucleic Acids Res 2015;43:5586–5600.

14. Bilinovich SM, Uhl KL, Lewis K et al. Integrated RNA Sequencing Reveals Epigenetic Impacts of Diesel Particulate Matter Exposure in Human Cerebral Organoids, Dev Neurosci 2020;42:195–207.

15. Amarasinghe SL, Su S, Dong X et al. Opportunities and challenges in long-read sequencing data analysis, Genome Biol 2020;21:30.

16. Wiener D, Schwartz S. The epitranscriptome beyond m(6)A, Nat Rev Genet 2021;22:119–131.

17. Zhou R, Xiao X, He P et al. SCAPE: a mixture model revealing single-cell polyadenylation diversity and cellular dynamics during cell differentiation and reprogramming, Nucleic Acids Res 2022;50:e66.

18. Yang J, McGovern A, Martin P et al. Analysis of chromatin organization and gene expression in T cells identifies functional genes for rheumatoid arthritis, Nat Commun 2020;11:4402.

